# When body mass index and listening to reasons for behavioral change make food choices healthier: Behavioral and neural mediators linking BMI and change interventions to healthier dietary decision making

**DOI:** 10.64898/2026.08.14.744895

**Authors:** Benjamin Flament, Belina Rodrigues, Iraj Khalid, Jean-Yves Rotgé, Christine Poitou, Hilke Plassmann, C Liane Schmidt

## Abstract

Resolving the inner conflict between improving one’s eating habits (change talk) and sticking to unhealthy ones (sustain talk) is a key target in communication-based behavioral change interventions such as motivational interviewing (MI). Recent work has shown that this inner conflict affects how tastiness and healthiness are traded off in dietary decision-making. The effect varied with body mass index (BMI). Here we aimed to identify why participants with higher BMI shifted toward healthier food choices after listening to change talk. An evidence accumulation model found that BMI affected health evidence sampling when listening to change talk, and taste evidence sampling when listening to sustain talk. A serial mediation analysis showed that the effect of BMI on change-talk-induced health evidence sampling was explained by stronger resting-state connectivity in the ventromedial prefrontal cortex within the default mode network (DMN), which in turn predicted greater motivation to change eating habits. These cross-sectional findings indicate that the intrinsic functional organization of valuation-related regions within the DMN is associated with motivation to change. They provide evidence that these neural and behavioral factors need to align with contextual cues, such as weight status (as reflected by BMI), and with reasons for behavioral change to promote healthier decision-making.

**Significance statement:** People often struggle to change unhealthy eating habits. Motivational interviewing (MI) addresses this challenge effectively in about 75% of cases (Rubak et al., 2005). The psychological and neural factors behind this success are not fully understood. We used brain imaging and computational modeling of food choices. After MI-generated change talk, participants gathered more information on food healthiness. Increased activity in the brain’s default mode network was observed in individuals with BMI ≥ 25 kg/m^2^ and predicted their motivation to change. This intrinsic brain activity-motivation link mediated the influence of weight status on health information gathering. These findings reveal that brain wiring interacts with personal motivation, explaining, in certain situations, like reflecting on why we want to change, how weight impacts choices.

## Introduction

Maintaining a healthy lifestyle and body weight is challenging and is often hindered by ambivalence about changing versus sustaining unhealthy eating habits (Kheniser et al., 2021). Communication-based interventions such as Motivational Interviewing (MI) aim to resolve this ambivalence (Miller and Rollnick, 2013). During MI, a patient verbalizes reasons to change unhealthy behaviors (change talk) and contrasts them with reasons holding them back (sustain talk).

We recently showed that during value-based dietary decision-making, listening to change talk led to healthier food choices and sustain talk to taste-based choices. These behavioral effects were most pronounced in participants with higher BMIs in the overweight and obesity range (BMI ≥ 25 kg/m^2^) (Rodrigues et al., 2026). Yet, it is unclear how change talk was particularly effective in helping these participants make healthier food choices. This study used a multipath mediation framework to investigate three main questions: (1) whether BMI is related to the brain’s intrinsic organization at rest, (2) if this resting-state activation serves as an inherent marker of personal motivation to change unhealthy habits, and (3) whether this sequential process mediates the effect of BMI on how evidence is accumulated during dietary decisions—especially when participants listen to their own statements about change.

In more detail, we modeled choices and reaction times with a drift-diffusion model (DDM) (Ratcliff et al., 2016) to gain access to the hidden variables of the dietary decision-making process. The model assumes that decision-making is a noisy process of evidence accumulation up to a threshold, at which a food item is accepted or rejected. DDMs are increasingly used to understand how people make choices (Roberts and Hutcherson, 2019). For example, how much they gather taste and health information during food choices (Maier et al., 2020; Sullivan and Huettel, 2021) or engage in self-regulation strategies (Ju et al., 2024). Comparing the model parameters across BMI groups and talk conditions can rule between alternative hypotheses: BMI and type of motivational talk might affect (1) how much a decision-maker sought healthiness vs. tastiness information about food during choice, (2) how biased the process is toward an option, or (3) how much evidence is needed to commit to an option. Based on our previous findings that change talk shifted food choices toward healthier over tastier options in participants with higher BMI (Rodrigues et al., 2026), we predicted that BMI should shape how strongly healthiness, relative to tastiness, drives evidence accumulation after change talk.

Turning to what might further drive these weight status effects in the decision process. Previous work has found that individuals with higher BMI exhibit altered, increased resting-state activation in the default mode network, linked to self-referential processing of food- and body-related information (Kullmann et al., 2012; Li et al., 2018). This may be reflected in the intrinsic functional organization of the brain’s default mode network (DMN), which is thought to engage during self-reflection on personal goals (Raichle et al., 2001; Buckner et al., 2008). A central hub of the DMN is the vmPFC, which is also a prominent part of the brain’s valuation system (Bartra et al., 2013; Clithero and Rangel, 2014), integrating long-term health goals and short-term taste goals during dietary decision-making (Plassmann et al., 2007, 2010; Hare et al., 2009, 2011; Hutcherson et al., 2012; Rudorf and Hare, 2014)(Plassmann et al., 2008; Hare et al., 2009, 2011; Hutcherson et al., 2012; Schmidt et al., 2018). Indeed, findings in participants with obesity show that resting-state vmPFC activity reconnects with another focal area of the brain’s valuation system, the ventral striatum, involved in motivational aspects of valuation when participants lost weight following bariatric surgery (Schmidt et al., 2021). In turn, such altered intrinsic functional organization may shape motivation to change, helping to explain why weight status affects dietary decision-making. Some studies have reported an association between higher BMI and greater motivation to change eating habits (Ljubičić et al., 2025). This greater motivation may reduce the health-taste conflict by increasing attention to health information during dietary decisions. The present study directly tested the combined effect of such behavioral and neural factors on healthier food choices across participants with varying BMI, who, as part of a broader research study, underwent MI to change their eating habits. We addressed this question by using computational modeling of the dietary decision-making process, resting-state functional magnetic resonance imaging (rest fMRI), and a multipath mediation framework. Moreover, we used a theory-driven approach and focused on the vmPFC within the DMN using an a priori defined region of interest reported in the literature for its key role during valuation (Bartra et al., 2013).

## Methods

### Ethics statement

The study was approved by the local ethics committees (CPP19.12.05 and CPP20.11.24.38405) and is part of the clinical trial C20-52 sponsored by INSERM. The study adhered strictly to the principles outlined in the Declaration of Helsinki. All participants provided written informed consent.

### Participants

Participants were recruited through public advertisements in the Paris area, France, and in collaboration with the Nutrition Department at Pitié-Salpêtrière Hospital in Paris. Participants were recruited based on the following inclusion criteria: age 18-70 years, right-handed, normal or corrected-to-normal vision, and ability to provide informed consent. The exclusion criteria consisted of substance abuse, neurological or psychiatric disorders, MRI contraindications (claustrophobia, non-removable metallic objects, pregnancy).

A total of 85 participants were initially recruited. A total of twenty participants were excluded from data analyses due to technical problems with the fMRI scanner and MRI contraindications (N=9), missing importance-to-change ratings (N=4), and movement outliers identified during the preprocessing of the resting-state fMRI data (N=7, framewise displacement > 0.5 mm or global BOLD signal changes > 3 standard deviations).

Thus, a total of 65 participants were included in analyses (52 women / 13 men; 32 with normal weight: BMI < 25 kg/m² / 7 with overweight: BMI between 25 - 30 kg/m² and 26 with obesity: BMI ≥ 30 kg/m², Table 1). The female-to-male ratio was comparable across all weight status groups.

**Table 1.** Participant characteristics by weight status groups

| Characteristics, means, [range] | Normal Weight Participants<br>BMI < 25 kg/m<sup>2</sup> | Overweight / Obese Participants<br>BMI ≥ 25 kg/m<sup>2</sup> |
| --- | --- | --- |
| Participants, n | 32 | 33 |
| BMI, kg/m<sup>2</sup> | 21.5 ± 2.1 / [17.0-24.9] | 36.6 ± 7.3 / [25.3-50.8] |
| Body Fat, % | 24.1 ± 6.7 / [10.8-38.6] | 45.2 ± 8.7 / [23.1-62.6] |
| Body Water, % | 53.2 ± 4.3 / [42.8-61.8] | 41.1 ± 6.1 / [32.5-53.7] |
| Age, years | 32.1 ± 12.2 / [21-58] | 37.1 ± 13.9 / [22-64] |
| Gender ratio, women:men | 27 : 5 | 25 : 8 |
| Education, years | 4.3 ± 1.7 / [0-8] | 3.0 ± 1.9 / [0-7] |

**Table 2.** Model fits

| Model | Conditions | DIC |
| --- | --- | --- |
| One drift weight : Health | Change Talk | 24079 |
|  | Sustain Talk | 24852 |
| One drift weight : Taste | Change Talk | 18873 |
|  | Sustain Talk | 17217 |
| Two drift weight : Taste and Health* | Change Talk (BMI $\leq 25$ kg/m<sup>2</sup>) | 8736 |
| | Change Talk (BMI $\geq 25$ kg/m<sup>2</sup>) | 8531 |
| | Sustain Talk (BMI $\leq 25$ kg/m<sup>2</sup>) | 8193 |
| | Sustain Talk (BMI $\geq 25$ kg/m<sup>2</sup>) | 8477 |

DIC – deviance information criteria; smaller DICs indicate better fit. \* – best-fitting model

Participants were classified as normal-weight when their BMI was below 25 kg/m² and as overweight or obese when their BMI was 25 kg/m² or higher. As shown in Table 1, the normal-weight group included participants with BMI values ranging from 17.0 to 24.9 kg/m², whereas the overweight/obese group included participants with BMI values ranging from 25.3 to 50.8 kg/m². Although two participants reported normal BMIs above 18.5 kg/m² at recruitment, their BMIs at the time of the MI session were 17 kg/m². Because their body-fat percentages remained within the healthy range (15% and 17%), they were retained in the analyses to avoid unnecessarily reducing the sample size.

### Experimental design

The experimental design shown in Figure 1 is described in detail by Rodrigues et al. 2026. In short, participants underwent a motivational interview (MI) session with a trained dietitian (Fig. 1b). At the end of the MI session, they responded to the question: "On a scale from 1 to 10, where 1 means ’Not at all’ and 10 means ’Very much’, how important is it for you to change your dietary habits?" This rating was used as a measure of motivation to change. It was assessed immediately following the MI session to elicit and strengthen motivation. Participants also rated another component of motivation, which was their readiness for change, using the same scale. However, the latter did not moderate vmPFC-DMN brain activation (even at a lenient, uncorrected, whole-brain threshold of p<0.001), nor did it moderate health-over-taste sampling during decision-making after change talk listening (r = 0.23, P = 0.064). We therefore focused our analyses on the specific importance-to-change component and refer to it as the motivation-to-change measure.

**Figure 1.**
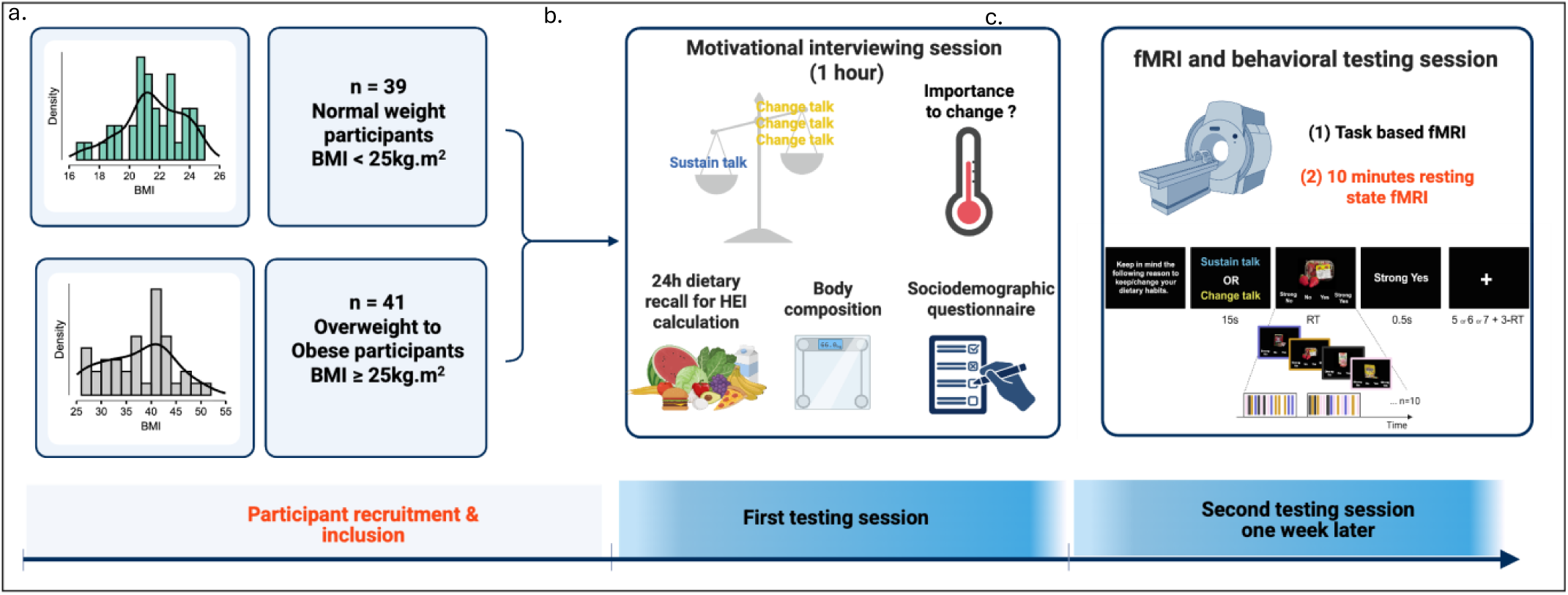
Study Design. Panels depict sequential events of the experimental design, which consisted of two visits separated by one week. After inclusion (a), participants with varying weight status (b) underwent a Motivational interviewing session during which they formulated reasons to change and sustain their eating habits and rated their motivation to change (importance to change rating) at the end of the session using a 10-point Likert scale. One week later (c), they underwent a task-based fMRI session reported in (Rodrigues et al., 2026) and a 10-minute resting-state functional magnetic resonance imaging (rest fMRI) assessment. This latter measure of the brain’s intrinsic functional organization at rest is the focus of this study.

One week after MI, participants underwent an fMRI experiment (see Fig. 1c) that included two task-based sequences reported by Rodrigues et al. 2026, and a 10-minute resting-state sequence (restfMRI). During restfMRI, participants were told to relax but remain awake for 10 minutes. They were further asked to look at a fixation cross displayed on the computer screen. None of the participants reported having fallen asleep when asked afterward.

Body composition (Body Mass Index (BMI), percent body fat, and water) was assessed after the MI session using a Tanita® Body Composition Analyzer SC-240MA (Tanita Corporation, Tokyo, Japan).

### Dietary decision-making task performed during fMRI

During the fMRI, participants performed a food choice task, the details of which are described in Rodrigues et al. 2026. Briefly, participants rated 150 food stimuli based on how much they wanted to eat each item at the end of the experiment. Before making these choices, they listened to recordings of their own change-and-sustain talk, which had been recorded during the MI session one week earlier. Choices were indicated on a 4-point Likert scale (from ’Strong No’ to ’Strong Yes’), and reaction times were recorded. Participants were informed that they would receive one food item, randomly selected from those they rated as ’Yes’ or ’Strong Yes’, to ensure an incentive-compatible experimental design.

## Statistical analyses

### Model-free analysis

Our dependent variables were participants’ food choices and their reaction times. These were analyzed using standard frequentist tests in JASP (version 0.19.3). Trials with reaction times below 200 ms were excluded from these analyses. Food wanting ratings were binarized into ’yes’ and ’no’ responses. This was done to parallel the DDM of binarized food choices outlined in the computational analysis section below, following the methods used by (Maier et al., 2020; Sullivan and Huettel, 2021; Khalid et al., 2024).

In more detail, to test whether the type of talk and weight status group (as reflected by BMI) impacted yes/no food choices and reaction times (RT), both dependent variables were analyzed using generalized linear mixed effect models (GLMM) for choices and linear mixed effects (LME) models for RTs. The models included fixed effects for talk, weight status group, and their interaction, as well as random intercepts nested by participant and random slopes for relevant fixed effects (see Equation (i).

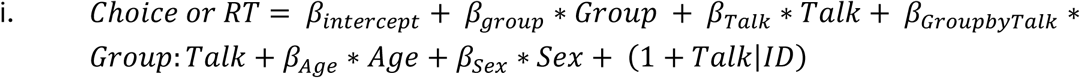

To account for the difference in distribution of the two dependent variables, the GLMM model was estimated using likelihood ratio tests (binomial distribution of choice), and the LME model was estimated using the Satterthwaite approximation (normal distribution of response time). Model fit was assessed using pseudo-R-squared for the GLMM (McFadden’s R² = 0.006) and variance-based R-squared for the LME (marginal R² = 0.024, conditional R² = 0.147) to determine the proportion of variance in the dependent variables explained by the models. Post hoc comparisons of average choices and reaction times between change and sustain talk were conducted using paired two-tailed tests.

### Model-based analyses

To examine the main questions of (1) whether listening to change versus sustain talk influenced the hidden variables of the decision-making process, and (2) whether this effect varied with weight status (i.e., BMI), the binarized food yes/no choices and corresponding reaction times were modeled by a hierarchical drift diffusion model (DDM).

#### Model Specification

The model was implemented in JAGS using the RWiener module extension (dwieners function) in RStudio, following similar approaches reported in (Khalid et al., 2024). The hierarchical structure of the DDM considered both group and individual differences in food decision-making. The model was fitted separately for each BMI group and talk condition, yielding four sets of DDM parameters (e.g., normal-weight group after change talk; normal-weight group after sustain talk; overweight and obese group after change talk; and overweight and obese group after sustain talk). In more detail, across trials of each of the four conditions, the dependent variable (Y) was the reaction time coded negatively for a "no" choice, and positively for a "yes" choice.

This dependent variable Y was then fitted by the DDM specified as follows:

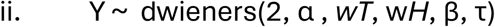

The boundary separation parameter (α) represented the threshold at which accumulation stopped and was capped at 2. The drift rate was scaled by the tastiness and healthiness ratings reflected by their respective weights (w_T_, w_H_). These two parameters quantified how much participants considered the healthiness and the tastiness information during evidence accumulation. The starting point bias parameter (β) reflected the initial propensity to accept or reject a food item. The non-decision time (τ) parameter represented the time required to start the evidence accumulation due to sensorimotor and psychomotor integration processes. Priors for each of these parameters are reported in Khalid et al. 2024.

#### Model estimation

The five parameters of the model (α, wT, wH, β, τ) were estimated using Gibbs sampling via Markov Chain Monte Carlo (MCMC) in JAGS to generate posterior values for each parameter. After 5,000 adaptation samples, three chains of 10,000 samples each were run with different starting values generated by three different random number generators. This resulted in a posterior chain of 30,000 plausible values per parameter, which were then used for comparisons across BMI groups and talk conditions.

Gelman-Rubin tests (Gelman and Rubin, 1992) confirmed convergence for each parameter, with potential scale reduction factors not exceeding 1.02 at participant or population levels. The deviance (log posterior) had a potential scale reduction factor ∼1.0. Autocorrelation between chains remained low across all groups and parameters (autocorrelation at lag 100 was approximately 0.0), indicating good chain performance.

Model selection

Three different versions of the generic DDM were compared: two single-weight models, which assumed that the drift rate was scaled by either tastiness (w_T_) or healthiness (w_H_) alone, and one two-weight DDM, which assumed that both sources of evidence scaled the drift rate.

To compare and select the best fitting model, the deviance information criteria (DIC) (Spiegelhalter et al., 2002) was calculated as follows :

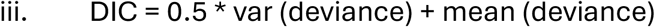

#### Parameter recovery

Parameter recovery for the drift diffusion model was assessed by fitting the winning hierarchical Wiener diffusion model, the two-weight DDM, to simulated datasets in R. Simulated datasets were created for each BMI group (Normal vs. OverweightCObese) and talk condition (Sustain vs. Change talk) using the sets of free parameters obtained from fitting the DDM to the observed dataset.

Parameter recovery was assessed by comparing posterior chains for each recovered parameter (*wT*, w*H*, β, τ, α) to posterior chains of the generating parameters in each of the four conditions (talk x BMI group). The recovery was quantified by the mean absolute deviation and Pearson’s correlation coefficient between generating and recovered parameters. Recovery was good to excellent with rates ranging between 90% and 92% for drift weights and 85% and 97% for noise, non-decision time, and starting point bias parameters. Thus, the winning DDM generated identifiable parameters.

#### Comparison of parameters between groups and talk conditions

To assess differences in parameters between BMI groups and talk conditions, the posterior value chains for each parameter were compared. In more detail, for each comparison (e.g., normal vs. overweight & obese BMI group, or change vs. sustain talk), posterior parameter values were subtracted from each other, leading to 30,000 differences per free parameter. These differences were then binarized as 1 for positive differences and 0 for zero or negative differences. The mean of these 30,000 binary values reflected the probability (ppMCMC) of a non-zero, positive difference, which should be greater than 95%, corresponding to a p_MCMC_ < 0.05.

#### Correlation of the model’s free parameters to motivation

To assess the link between participants’ motivation, the health-versus-taste DDM parameter difference was calculated by subtracting the drift-rate for tastiness (w_T_) from that for healthiness (w_H_). This score ranged from -1 to +1. Negative values indicated a greater influence of tastiness on the drift rate, controlling for healthiness, while positive values showed a greater influence of healthiness, controlling for tastiness. In other words, when the health-versus-taste difference was positive, the participant’s choice was more shifted toward sampling healthiness taste evidence.

Correlations between motivation to change and this DDM parameter difference were assessed using Pearson’s correlations, and differences in the Pearson’s correlation coefficients between types of talks and BMI groups were evaluated with Fisher’s r-to-z transformation tests (Cohen, 2013).

Furthermore, we tested the robustness of detected correlations by conducting a leave-one-sample-out (LOSO) cross-validation. To this aim, the dataset was divided into 65 training sets, each containing 64 participants, with one participant left out as the test set. For each training set, a general linear model (GLM) was performed to obtain a beta coefficient using the MATLAB glmfit function. This coefficient, along with the test set’s x variable (i.e., residuals with age and gender confounds regressed out), was used to predict the z-scored y using the glmval function in MATLAB. The process was repeated 65 times, resulting in 65 predictions (yhats). These predicted values were subsequently regressed against the actual observed, z-scored y values using the fitlme function in MATLAB. Prediction accuracy was assessed with MSE, RMSE, and R-squared metrics, indicating how accurately the predicted variable (yhats) matched the observed one.

## Resting state fMRI acquisition

Resting-state fMRI was measured with a Verio 3T Siemens MRI scanner. The resting-state session lasted 10 minutes and was conducted following the food choice task and a Multi-Source Interference task.

Multi-echo echo-planar images (mEPIs) were acquired with the following sequence: Three echo times of 14.8 ms, 33.4 ms, and 52.1 ms, repetition time (TR) = 1.25 s, flip angle = 68°, in-plane resolution = 3.0 × 3.0 mm², slice thickness = 3.0 mm (no gap), 48 axial slices, field of view = 210 mm, acquisition matrix = 70 × 70.

## MRI Data Analysis

Analyses of fMRI data were performed using Statistical Parametric mapping (SPM12, RRID:SCR_007037, release 12.7771) and the Connectivity toolbox (CONN RRID:SCR_009550, release 22.v2407) (Whitfield-Gabrieli and Nieto-Castanon, 2012).

### Preprocessing

Functional and anatomical images were preprocessed in SPM 12 and the CONN toolbox. First, at each TR, the three mEPIs were summed (Gowland and Bowtell, 2007; Poser and Norris, 2009; Kettinger et al., 2016). In SPM12, the summed EPIs were then realigned and unwarped to correct for head motion and susceptibility-related distortions, co-registered to the first functional volume, and resampled using b-spline interpolation. Slice-timing correction was applied to adjust for differences in acquisition time across slices. Volumes with excessive motion or global signal change were flagged as outliers (framewise displacement > 0.5 mm or global BOLD signal change > 3 SD) and later included in denoising and statistical analyses as nuisance control regressors. Structural and functional images were normalized to Montreal Neurological Institute (MNI) space using unified segmentation parameters from the anatomical MRI, which was segmented into grey matter, white matter, and cerebrospinal fluid, and resampled to 2 mm isotropic resolution. Finally, realigned, unwarped, slice-time-corrected, and normalized EPIs were spatially smoothed with an 8- mm FWHM Gaussian kernel.

### Denoising

Denoising of these SPM12 preprocessed images was conducted in the CONN toolbox. Nuisance regressors included principal components from white matter (10 components) and CSF (5 components) signals (CompCor) (Behzadi et al., 2007), six motion parameters and their first derivatives, the identified outlier volumes, session effects and their first derivatives, and linear trends. Residual BOLD time series were then bandpass filtered between 0.008 and 0.09 Hz to reduce low-frequency drifts and high-frequency noise.

### Group independent component analysis (ICA)

To identify large-scale functional networks, a group independent component analysis (ICA) was performed on the preprocessed, denoised resting-state brain images. Subject-level dimensionality reduction was achieved using singular value decomposition, followed by a second reduction step at the group level. A fastICA algorithm was then applied to estimate eight spatially independent, temporally coherent components across all participants, and subject-specific maps for each component were obtained using back-projection (Hyvarinen, 1999; Calhoun et al., 2001).

### Regions of interest (ROI)

The region of interest in this study was localized within the ventromedial prefrontal cortex (vmPFC). It was defined by the Montreal Neurological Institute (MNI) coordinates [-2, 40, -8], reported by (Bartra et al., 2013) for vmPFC activation in response to subjective value at the time of choice. A 5 mm sphere centered at MNI coordinates [-2, 40, -8] was used for small-volume correction of Statistical Parametric Maps (SPMs) and for extracting beta values reflecting vmPFC activation within the DMN identified by ICA for mediation analyses.

## Univariate statistical analyses of brain resting-state data

Group-level analyses of DMN activity strength derived from independent component analysis conducted within the general linear model framework implemented in CONN/SPM. For each voxel, GLMs tested the effects of group and other subject-level predictors on first-level connectivity measures such as continuous BMI values, Importance-to-change ratings, and health-versus-taste evidence sampling. Statistical inference was performed at the cluster level using Gaussian random field theory (Worsley et al., 1996; Nieto-Castanon, 2020), with a cluster-forming threshold of p < 0.001 (uncorrected) at the voxel level and a cluster-level p-FDR < 0.05 to correct for multiple comparisons.

## Multivariate mediation analysis

A cross-sectional (i.e., single-level), serial mediation analysis was conducted using the Multilevel Mediation and Moderation (M3) toolbox (Wager et al., 2008) implemented in MATLAB. The serial mediation tests the hypothesis that an independent variable (X) influences a dependent variable (Y) through a sequential chain of two mediators (M1 → M2), where the effect flows from X to M1, from M1 to M2, and from M2 to Y. All variables were z-scored (standardized to mean = 0, SD = 1) prior to analysis to facilitate comparison of path coefficients across different measurement scales.

More specifically, the mediation model estimated the following pathways: Path b₁ represented the effect of BMI (X) on the vmPFC resting state activity within the DMN (M1), while path b₂ captured the effect of vmPFC activity within the DMN (M1) on the importance to change ratings (M2), controlling for the effect of BMI (X). Path b_3_ estimated the effect of importance to change ratings (M2) on the health-versus-taste information sampling during dietary decision-making (Y), controlling for the effects of BMI (X) and vmPFC activity within the DMN (M1).

The total effect of BMI (X) on health-versus-taste drift rate weights of the DDM (Y) was represented by path c, while the direct effect between these two variables after accounting for both mediators was represented by path c’ coefficients.

Mediation was inferred from the serial indirect effect, computed by the following equation:

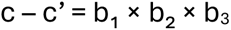

Bootstrap tests (10,000 iterations) were used for significance testing of each path’s coefficients.

It is worth noting that the importance of change ratings (M2) was measured immediately after the MI session, and one week prior to the fMRI session at which the DMN resting-state activity (M1) and the health-versus-taste weighting (Y) were measured. This was a deliberate choice because we wanted to leverage the MI session to motivate personal change. The cross-sectional design of the mediation analysis examines tendencies across different participants rather than within the same individuals, which mitigates the need to assume a specific order of cause and effect.

Moreover, for completeness, we checked for the effect of alternative orderings of the mediators, such as motivation (M1) -> vmPFC (M2) -> health-versus-taste drift weights (Y). This ordering did not lead to a significant indirect effect of BMI (X) on health-versus-taste sampling after change talk (Y) (path b1*b2*b3: β = 0.04, p = 0.135).

## Sensitivity analysis

Body Mass Index (BMI) is an anthropometric measure and does not define metabolic health (Sommer et al., 2020). Recent obesity frameworks distinguish BMI from adiposity and clinical status ((Rubino et al., 2025; Busetto et al., 2024). The body composition assessment of participants after the MI session also provided estimates of body fat percentage. In a sensitivity analysis, we tested whether this alternative measure of weight status was associated with (a) motivation to change and (b) health-versus-taste DDM drift rate weights.

## Results

### Behavioral results

The observed behavioral effects are described in detail by Rodrigues et al. 2026. Briefly, all participants chose to reject food more after listening to change talk compared with sustain talk (χ² = 4.50, p = 0.034, Fig 2a), with no effect of type of talk on reaction times (F = 0.211, p = 0.648, Fig 2b). Participants with overweight or obesity rejected food more (χ² = 8.07, p = 0.005, Fig. 2a) and chose faster (F = 13.97, p < 0.001) than normal-weight participants (< 25 kg/m^2^). When modeling these effects with a DDM to examine behavioral differences in the hidden decision-making process, we found that the drift rate of evidence accumulation was more driven by healthiness than by tastiness among participants with overweight or obesity (p_MCMC_ = 0.02). This difference between weight status groups was further amplified when considering only change-talk trials, relative to the sustain talk conditions (p_MCMC_ = 0.004; Fig. 2d). In addition, participants with obesity and overweight also had shorter non-decision times (τ) than normal-weight participants (p_mcmc_ = 0.03), suggesting faster sensory and psychomotor processing to accept or reject a food.

**Figure 2.**
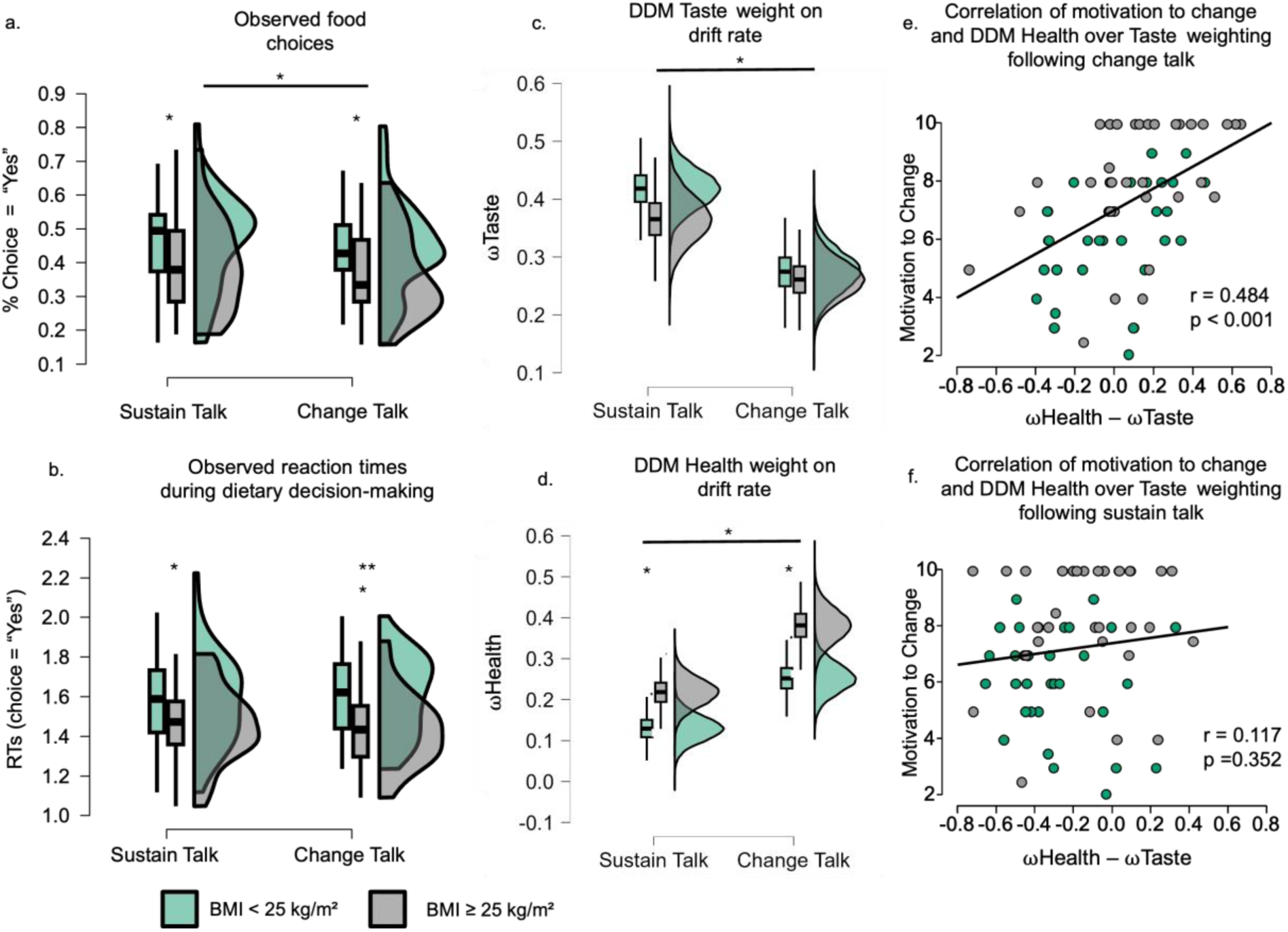
Behavioral and Computational results. All boxplots display 95% confidence intervals, with boxes indicating the interquartile range from the 25th percentile (Q1) to the 75th percentile (Q3). The black lines indicate medians, and the whiskers extend from the minimum to the maximum values. In green are the normal-weight participants with a BMI < 25kg/m², and in grey are the overweight and obese participants with a BMI ≥ 25kg/m². **(a)** Observed food choices (% of ‘yes’), and **(b)** reaction times (RT). **(c and d)** DDM drift-rate weight parameters for tastiness and healthiness of food items at choice. **(e and f)** Correlation of motivation to change (“How important is it to change your eating habits?”) with the difference in health and taste evidence sampling (i.e., drift-rate weight DDM parameters) after (e) change and (f) sustain talks. r – Pearson’s correlation coefficient. * p < 0.05, *** p < 0.001, two-sample and paired two-tailed t-tests.

Given these observed and modeled behavioral results, we examined our main question: Why do participants with overweight or obesity prioritize health over taste during evidence accumulation?

We tested (1) the association with motivation to change, investigated if such associations would be (2) driven by differences in intrinsic brain patterns at rest, and (3) linked them within the framework of a serial mediation.

#### (1) Association of BMI and DDM parameters with motivation to change

Self-reported motivation to change correlated significantly with the health-versus-taste drift rate weight difference following change talk (r = 0.484, p < 0.001, fig 2e), whereas no significant correlation was observed for sustain talk (r = 0.117, p = 0.352, fig 2f). The significant association between motivation to change and healthiness sampling during decision-making after change talk also generalized. The LOSO cross-validation (β = 0.18, t(2,63) = 3.6, p = 0.0005, 95%CI [0.08; 0.29], R² = 0.64, MSE = 0.08, RMSE = 0.29) indicated that the theoretically derived motivation calculated by observed healthiness sampling from the training set predicted test out-of-sample, observed motivation to change with 64% accuracy.

Taken together, these results indicated that participants who were more motivated to change sampled more evidence about the healthiness of foods and less about their tastiness during dietary decision-making.

Moreover, motivation to change was also significantly correlated with weight status as approximated by BMI. The higher the BMI, the greater the motivation to change (r = 0.51, p < 0.001). The association between weight status and motivation to change also held when weight status was defined as percent body fat (r = 0.45, p < 0.001). Moreover, it was assessed via a LOSO cross-validation, which was significant (β = 0.11, t(2,63) = 2.58, p = 0.012, 95% CI [0.03; 0.21], R² = 0.58, MSE = 0.06, RMSE = 0.25). The association was driven by participants with overweight or obesity (BMI> 25 kg/m²; r = 0.40, p = 0.019). It was not significant for normal-weight participants (r = −0.30, p = 0.09). The between-group difference in correlation was significant by Fisher’s r-to-z transformation (z = 2.85; two-tailed p = 0.0044).

#### (2) Differences in intrinsic brain patterns at rest

Resting-state functional connectivity analysis revealed a significant correlation of intrinsic vmPFC activation within the DMN to motivation to change ratings (MNI = [-4, 40, -4], small- volume corrected (SVC) p_FWE_ = 0.019, family-wise error correction, Fig. 3a), and a health- versus-taste DDM parameter difference (MNI = [0, 40, 0], SVC p_FDR_ = 0.034, false discovery rate corrected, Fig 3b).

**Figure 3.**
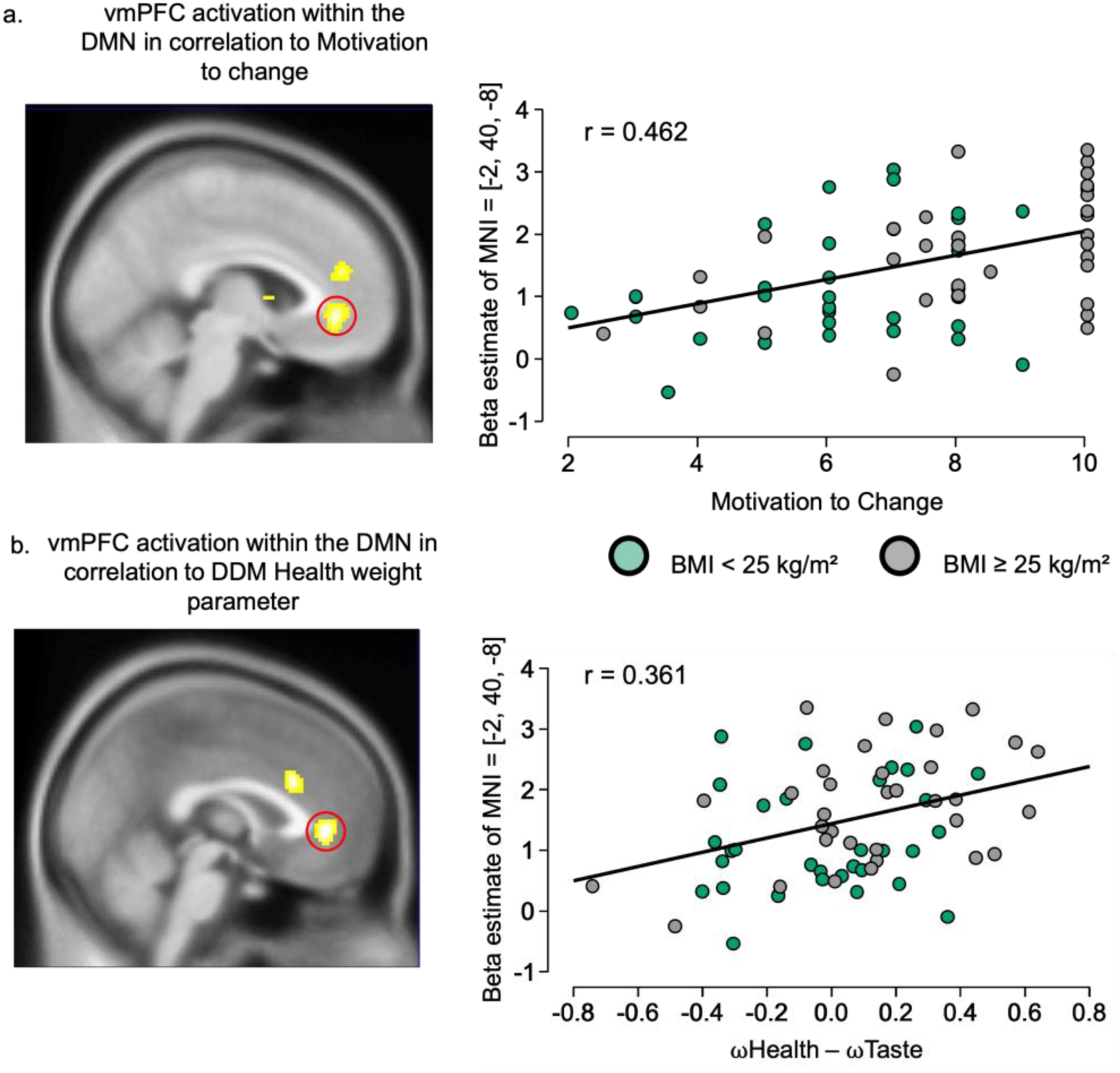
Correlation of vmPFC resting state activation with motivation to change and DDM parameters. (a) Statistical parametric maps (SPM) show yellow voxels that significantly correlated with motivation to change ratings (small-volume-corrected p_FWE_ < 0.05), and (b) DDM health-versus-taste drift rate weight difference (small-volume-corrected p_FDR_ < 0.05). These are overlaid on the average structural brain image and displayed at a whole-brain p < 0.001 uncorrected threshold. The scatter plots display the correlations between resting-state vmPFC activity, extracted from an a priori defined ROI (MNI=[-2, 40, - 8], reported by Bartra et al. 2013), and motivation to change ratings, as well as the health- versus-taste drift rate weight difference from the DDM fitted to food choices made when listening to change talk. Each dot represents an individual participant, and r values indicate Pearson’s correlation coefficients. In green are the normal-weight participants with a BMI < 25kg/m², and in grey are the overweight and obese participants with a BMI ≥ 25kg/m².

**Figure 4.**
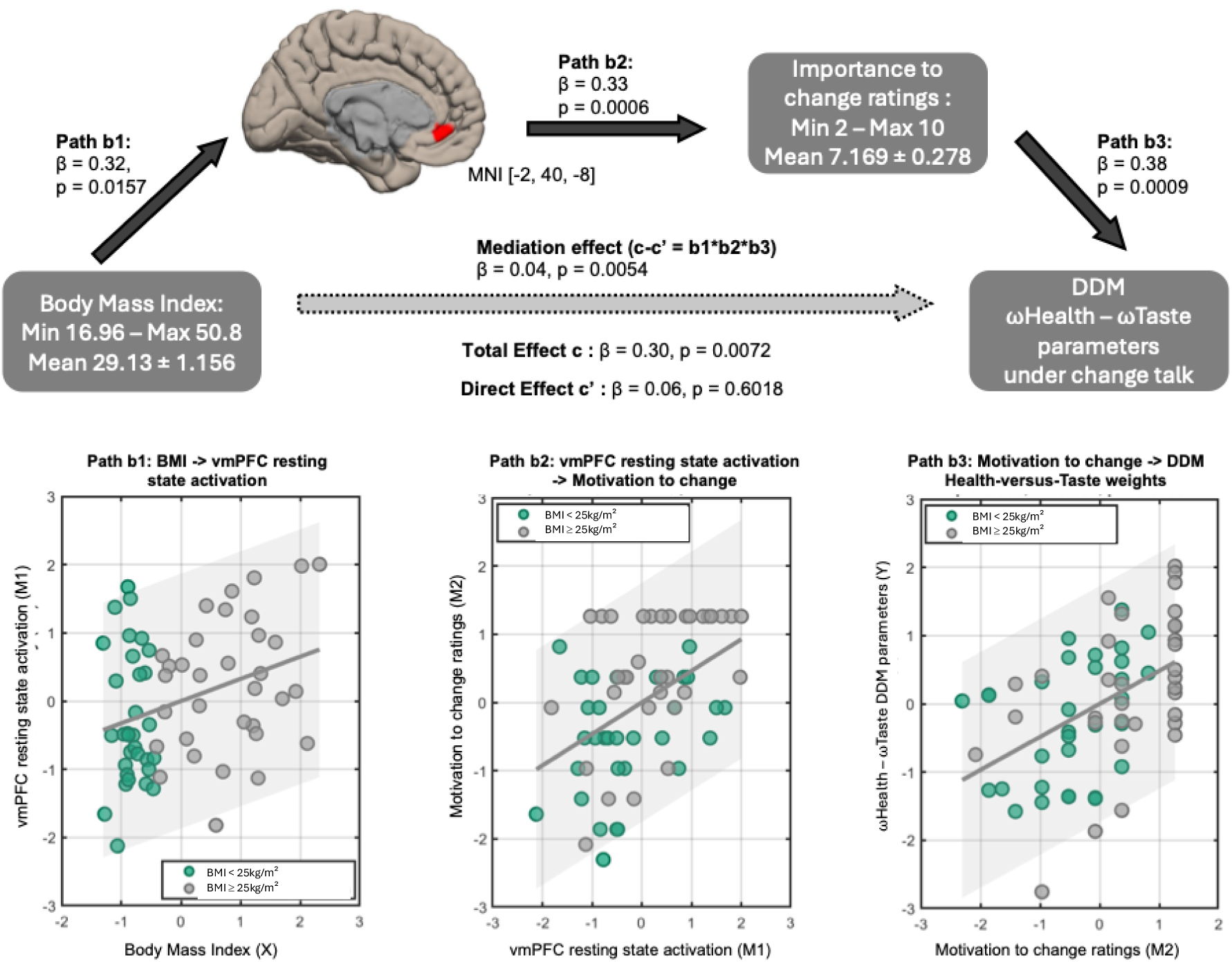
Three-path mediation results. The vmPFC ROI is centered at MNI = [−2, 40, −8], as reported by Bartra et al., 2013. Path b1 shows the effect of BMI on vmPFC activity within the DMN. Path b2 represents the effect of vmPFC activity within the DMN on motivation-to- change ratings, controlling for BMI. Path b3 illustrates the effect of motivation-to-change ratings on the relative weight of healthiness over tastiness of food items on the drift rate of the DDM model of dietary decision-making. Scatterplots display the correlations between variables for each regression path, with dots representing individual participants. In green are the normal-weight participants with a BMI < 25kg/m², and in grey are the overweight and obese participants with a BMI ≥ 25kg/m².

As shown in Figure 3a and b, participants who were more motivated to change their dietary habits (Pearson’s r = 0.46) and had a stronger health-versus-taste drift rate weight (Pearson’s r = 0.36) displayed stronger resting-state vmPFC activation within the DMN. The correlation between vmPFC resting-state activation and the health-versus-taste drift rate weight was specific to dietary decision-making trials following change talk. This correlation was non-significant for decision-making trials following sustained talk.

Moreover, the associations between vmPFC DMN activation to motivation and healthiness sampling after change talk generalized as indicated by LOSO cross-validation (motivation: β = 0.09, t(2,63) = 2.25, p = 0.027, 95%CI [0.01; 0.17], R² = 0.57, MSE = 0.05, RMSE = 0.23; health-vs-taste drift weight: β = 0.10, t(2,63) = 2.44, p = 0.017, 95%CI [0.02; 0.19], R² = 0.57, MSE = 0.06, RMSE = 0.24).

In summary, these correlational findings reveal that participants with stronger vmPFC activity at rest reported greater motivation to improve their diet and displayed more health information sampling during dietary decision-making.

#### (3) Linking vmPFC activation within the DMN at rest to motivation to change for explaining weight status effects on health-over-taste advantage during dietary decision making after change talk

To identify putative mediators of the association between BMI and the healthiness-over- tastiness sampling during decision-making (total effect c: β = 0.30, p = 0.0072), a serial mediation analysis was conducted.

BMI significantly predicted vmPFC resting-state activation (path b1: β = 0.32, p = 0.0157). Participants with higher BMIs exhibited greater vmPFC resting-state activation within the DMN. The vmPFC resting-state activation then significantly predicted greater motivation to change current eating habits (path b2: β = 0.33, p = 0.0006). Finally, motivation to change significantly predicted increased health-versus-taste sampling during dietary decision-making (path b3: β = 0.38, p = 0.0009). Taken together, these three path regressions significantly reduced the direct effect of BMI on health-versus-taste sampling (direct path c’: β = 0.06, p = 0.6), indicating a full indirect effect via vmPFC resting-state activation and motivation to change (indirect, mediation path b1*b2*b3: β = 0.04, p = 0.0054).

A sensitivity analysis showed that another measure of weight status, percent body fat, did not yield a significant total effect on the health-versus-taste drift-rate weights (path c: β = 0.22, p = 0.06). Although this association was borderline non-significant, it was positive, indicating that participants with greater body fat accumulated more information about the healthiness of food during dietary decision-making after change talk statements.

Taken together, these results showed that the association between weight status, specifically its definition by BMI, and health information sampling during food decision-making is statistically mediated by resting-state vmPFC activation within the DMN and the motivation to change eating habits.

## Discussion

In this study, we investigated why people with higher BMI prioritized health over taste in dietary decision-making, particularly after hearing their change talk statements from MI. We found that participants with higher BMI had stronger vmPFC activation within the DMN and were more motivated to change. A three-path mediation analysis formalized that BMI influenced health-based food choices indirectly through vmPFC resting-state activation and motivation to change current eating habits.

We used a mathematical model to formalize observed food choices, thereby allowing us to isolate specific latent psychological subprocesses that behavioral measures alone could not disentangle. DDMs have demonstrated sensitivity to a range of influences on dietary decisions, such as hunger states (March and Gluth, 2025), beliefs about hunger (Khalid et al., 2024), and tendencies to select default choice options (Sullivan et al., 2025). The model revealed that participants with overweight and obesity exhibited faster psycho- and sensorimotor integration, in line with faster reaction times, and, especially after listening to change talk, shifted toward accumulating more evidence from health information. This finding stands in contrast to research showing that health considerations in dietary decision-making often receive fewer attentional resources, or if they do, take more time (Hutcherson et al., 2012; Maier et al., 2020; Sullivan and Huettel, 2021).

Our cross-sectional mediation results can explain this divergence. We employed a theory-driven approach and focused on the mediating role of a specific functional brain organization, namely the intrinsic activation of the vmPFC within the DMN identified using ICA (Raichle et al., 2001). Under task-based fMRI, the vmPFC is known to play a central role in value-based decision-making (Bartra et al., 2013). At rest, it coactivates with other brain regions to form the DMN, a prominent brain network that activates more at rest in participants who engage more in self-referential thinking, recall personal experiences, or envision future scenarios (Cabeza and St Jacques, 2007; Schacter et al., 2007; Benoit et al., 2014). Weight status has been shown to be associated with functional resting–state connectivity of the vmPFC to a motivational hub – the ventral striatum (Schmidt et al., 2021). Our results consolidate these different lines of research by providing novel evidence that participants who activated the vmPFC within the DMN more often were also more motivated. This association then fully mediated the effects of weight status on health-information sampling during dietary decision-making. The specificity of this finding to the contextual condition of listening to change talk further suggests that the associations between weight status, increased vmPFC resting-state activity within the DMN, and motivation to change alone are insufficient. These effects require contextual cues such as listening to statements that favor behavioral change to translate into healthier dietary decision-making.

A potential confound in the resting-state fMRI results could be a carryover effect from the prior food-choice task. However, since more than 10 minutes elapsed after the food choice task—during which time participants engaged in another task measuring interference resolution—the likelihood of a carryover effect is small. Indeed, task-induced changes in functional connectivity have been shown to dissipate within approximately five minutes of task cessation (Tung et al., 2013), and the dominant resting-state connectivity pattern of the default network has been characterized as temporally stable and insensitive to preceding task states (Grigg and Grady, 2010). When carry-over effects do persist, they tend to concentrate in task-positive network regions such as the supplementary motor area and precentral gyrus, unlike default network activity, which is inherently oriented toward internal cognition (Grigg and Grady, 2010). The elevated vmPFC connectivity observed in this study may therefore reflect ongoing self-referential processing relevant to health motivation, rather than residual noise from the preceding tasks. Future studies should employ longitudinal designs that track resting-state brain activity, motivation, and decision-making parameters throughout an actual weight-loss journey or over longer periods to determine whether the mediation pathways we identified are malleable.

Participants with obesity were recruited from a hospital setting at the start of a weight loss treatment course. This recruitment strategy may be an important confound because treatment-seeking status may itself be associated with importance to change, as patients with obesity have been shown to be more concerned about the consequences of unhealthy eating habits (Tang et al., 2012). Such heightened concern could indicate that they are less ambivalent about changing their eating habits (Koehler and Leonhaeuser, 2008), which could translate into more motivation to change and challenge a deficit-oriented view of overweight and obesity. Moreover, participants seeking obesity treatment may have experienced greater exposure to weight-related stigma or repeated pressure to change their weight and eating behaviors (Puhl and Heuer, 2009). Such experiences can be internalized and may influence self-regulation, motivation, and food-related decision-making (Pearl and Puhl, 2018). Weight-related social identity threat may therefore represent an additional unmeasured factor contributing to the association between BMI and importance to change observed here. Future studies should assess experienced and internalized weight stigma to test this possibility. However, according to the World Health Organization, the primary challenge patients with overweight and obesity face is not a lack of motivation to change but rather translating that motivation into weight loss amidst obesogenic environments, socio- economic pressures, and metabolic alterations (Organization, 2018). Here, we show that contextual cues, such as hearing the reasons for change prior to a food choice, can play a crucial role.

All of the correlational results were statistically generalizable using leave-one-out cross-validation. Future studies should further test the generalizability of these findings outside the experimental, laboratory setting by recruiting participants with obesity who are not seeking medical weight-loss treatment.

Motivation to change was assessed after the MI session by asking participants to rate how important change was to them. MI research suggests that motivation involves more than just recognizing the importance of changing unhealthy behaviors (Mason et al., 2010). It also includes other components, such as readiness to change. However, our results showed that readiness ratings did not yield significant outcomes, and thus primarily reflect how important participants view behavioral change for themselves.

Looking ahead, our study provides preliminary evidence from a laboratory experimental setting that could inform the development of future adaptive interventions to change actual food intake, long-term dietary adherence, or weight. For instance, future studies could test the hypothesis that incorporating change-talk reminders into daily routines, perhaps by using the person’s own voice through digital tools and applications, can support healthier real-world food choices. Moreover, research has shown that replaying autobiographical memories, engaging in mental imagery of one’s future healthier self, or mindfulness training activates and strengthens the DMN at rest (Taylor et al., 2013; Garrison et al., 2015; Rahrig et al., 2022). The association between vmPFC-DMN activity and importance to change suggests that these techniques could be tested in future longitudinal studies to determine whether they improve real-world dietary behavior or adherence. Lastly, our computational results propose that ambivalence toward adopting healthier eating habits could be resolved by structuring MI conversations to prompt an attentional shift from taste to health focus in therapeutic settings.

In conclusion, our findings contribute to a broader theoretical understanding of behavior change and the intention-behavior gap, in which individuals are motivated to change, but do not consistently adhere to healthier diets over the long term. Our findings suggest that successful dietary change requires alignment across multiple levels: intrinsic functional brain organization associated with motivation to change, and contextual cues that activate that motivation. When these factors align, healthier choices are more likely. They provide insights that contribute to the translation into personalized interventions to support healthier, real-life dietary behavior and improve clinical outcomes.

## Conflict of interest statement

The authors declare no competing financial interests.

## Acknowledgments

This project has received funding from the Foundation NRJ — Institut de France. BR received a fellowship of the Paris Region Fellowship Program supported by the Paris Region, which received funding from the European Union’s Horizon 2020 research and innovation program under the Marie Skłodowska-Curie. The funders had no role in study design, data analysis, manuscript preparation, or publication decisions. This study is part of the clinical trial C20-52 sponsored by Inserm.

## Notes

### Competing Interest Statement

The authors have declared no competing interest.

### Summary of Updates

The manuscript terminology was refined to improve clarity and accuracy. The Methods section was revised, including the addition of a table summarizing participant characteristics. Minor changes were also made throughout the manuscript in response to co-author feedback.

